# AF2Rank Revisited: Reproducing AlphaFold-Based Structure Evaluation and a Hypothesis for Context-Aware Refinement (CAR-AF)

**DOI:** 10.1101/2025.04.30.651434

**Authors:** Priyanshu Kumar

## Abstract

Protein structure prediction has undergone transformative advancements with AlphaFold2 achieving near-experimental accuracy across extensive protein datasets. This study reproduces and validates the AF2Rank pipeline, which utilizes AlphaFold’s intrinsic confidence metrics—predicted Local Distance Difference Test (pLDDT) and predicted Template Modeling score (pTM)—to evaluate and rank decoy protein structures without dependence on multiple sequence alignments (MSAs). The pipeline was implemented locally using the Rosetta decoy dataset, overcoming reproducibility challenges such as software dependency conflicts, residue indexing inconsistencies, and system-level execution issues. This framework successfully enabled high-confidence evaluation of over 1000 decoys for a benchmark target (1a32), with ongoing expansion to 133 protein targets.

Notably, we discovered that AlphaFold confidence metrics encode protein-specific “fingerprints,” enabling reverse classification of structures to their source proteins. Our XGBoost-based classifier achieved 61.5% accuracy across 133 distinct protein targets, substantially exceeding random chance (0.75%). Feature importance analysis revealed that traditional energy functions (Rosetta normalized score, 26%) and derived interaction features (pLDDT-pTM ratio, 16%) provide the strongest discriminative power, suggesting AlphaFold metrics capture meaningful biological and structural information beyond generic quality assessment. Building upon this foundation, a novel hypothesis is introduced: Context-Aware Refinement of AlphaFold Predictions (CAR-AF). This approach postulates that AlphaFold-predicted structures may be further improved through refinement in the presence of their native binding partners—such as receptors or ligands—thereby producing conformations that are both structurally and functionally enhanced. The proposed methodology comprises four stages: generation of initial predictions via AlphaFold, structural modeling of relevant binding partners, docking of predicted proteins into these biological contexts, and refinement of the resulting complexes using molecular modeling tools such as Rosetta or HADDOCK. Structural and energetic metrics including TM-score, RMSD, and binding energy will be used to assess potential improvements in predictive quality.

## I. Introduction

Proteins serve as the molecular workhorses of life, executing essential functions such as catalysis, transport, and structural support. Accurately determining a protein’s three-dimensional conformation is crucial for understanding its biological role and guiding rational design in drug discovery, enzyme engineering, and other biotechnological applications. Historically, experimental methods like X-ray crystallography or NMR spectroscopy were the primary means of resolving protein structures, but these approaches can be labor-intensive, expensive, and limited in throughput.

**Fig. 1.**
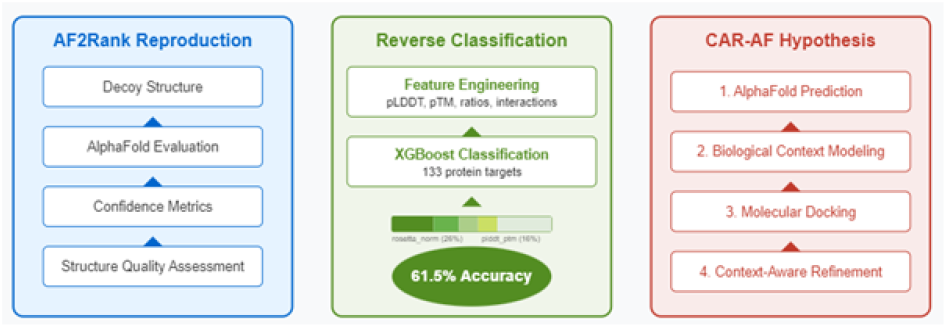
Graphical Abstract of AF2Rank Reproduction and CAR-AF Hypothesis. This figure illustrates the three key components of our research. Left panel: The AF2Rank pipeline reproduction, which processes protein decoy structures through AlphaFold2 evaluation to generate confident structure quality assessments without multiple sequence alignments. Middle panel: Our novel reverse classification approach, achieving 61.5% accuracy in identifying protein targets across 133 classes based solely on AlphaFold confidence metrics, with rosetta_norm (26%) and plddt_ptm_ratio (16%) as the most discriminative features. Right panel: The proposed Context-Aware Refinement (CAR-AF) methodology, comprising four sequential stages: AlphaFold prediction, biological context modeling, molecular docking, and context-aware refinement. Bottom panel: Conceptual illustration of the CAR-AF process, from isolated structure prediction through complex formation to context-refined structure, demonstrating how incorporating biological binding partners can enhance both structural accuracy and functional relevance of protein models.

Recent innovations in machine learning, most notably AlphaFold2 [1], have revolutionized computational protein structure prediction by providing near-experimental accuracy for a wide range of proteins. AlphaFold2 achieved unprecedented performance at CASP14 (Critical Assessment of protein Structure Prediction) [15], with a median GDT (Global Distance Test) score of 92.4 across targets. While AlphaFold offers confidence metrics such as predicted Local Distance Difference Test (pLDDT) and predicted Template Modeling score (pTM) to judge the reliability of its models, the full potential of these scores remains only partially explored.

The AF2Rank pipeline, proposed by Jing et al. [2], demonstrated that AlphaFold’s internal scoring mechanisms could be used to compare and rank decoy structures without reliance on multiple sequence alignments (MSAs) or extensive computational resources. This approach offers significant advantages for high-throughput structure evaluation, particularly when dealing with novel protein designs or sequences with limited evolutionary information.

This study addresses two primary objectives: (1) to reproduce and validate the AF2Rank methodology in a local computing environment, and (2) to propose a novel hypothesis for Context-Aware Refinement of AlphaFold Predictions (CAR-AF). The AF2Rank reproduction involved overcoming numerous technical challenges, including dependency conflicts, system compatibility issues, and data formatting constraints. Upon successful implementation, the pipeline was applied to evaluate protein structure decoys from the Rosetta dataset.

The CAR-AF hypothesis represents a conceptual extension beyond isolated structure prediction. It posits that AlphaFold predictions, while highly accurate in isolation, might benefit from refinement within their biological context—such as in the presence of binding partners, ligands, or receptors. This approach acknowledges that proteins in biological systems rarely function in isolation, but rather adapt their conformations in response to molecular interactions [13]. By integrating contextual information into the refinement process, the CAR-AF methodology aims to bridge the gap between geometric accuracy and functional relevance in protein structure prediction.

## II. Background and Motivation

The number of sequenced proteins is growing at a staggering pace. Public repositories such as UniProt contain roughly 220 million protein entries, yet only about 180,000 of these have experimentally determined structures [3]. This immense gap between sequence data and available structural information poses a critical challenge in molecular biology and drug discovery. Although methods like X-ray crystallography, Nuclear Magnetic Resonance (NMR), and cryoelectron microscopy have provided high-resolution insight into many structures, they are notoriously time-consuming, costly, and often fail for large or flexible proteins. A typical crystallography project may require 6-18 months of dedicated effort, with success rates as low as 30% for novel proteins. Similarly, high-resolution cryo-EM studies might cost upwards of $100,000 per structure and require specialized equipment with lengthy wait times for access. Accelerating structure determination is thus a top priority: beyond its scientific value, a fast and reliable pipeline could inspire solutions to widespread problems ranging from rising global temperatures to the urgent need for sustainable waste management.

While single-protein structure prediction has historically received the most attention, many essential biological functions depend on multi-chain protein complexes. Approximately 65% of proteins in eukaryotic cells function as part of complexes rather than as isolated entities. Accurately predicting these quaternary structures remains an active frontier. Traditional template-based and free-docking approaches often struggle when homologous templates are sparse or when the complex contains flexible regions. Success rates for complex prediction using conventional docking methods rarely exceed 40%

Much of the recent progress owes its origins to advances in coevolutionary analysis, where correlated mutations within MSAs highlighted physical proximity or inter-residue interactions [10]. This approach laid the groundwork for neural-network-based pipelines by identifying which amino acids likely contact one another in 3D space. Direct Coupling Analysis (DCA) methods, which can detect correlated evolutionary patterns across distant sites in sequences, achieved early successes with contact prediction accuracy rates of 70-80% for the top L/5 predicted contacts (where L is the sequence length). From there, models such as RoseTTAFold, OpenFold, and especially AlphaFold2 used deeper architectures (often transformer-based) and more sophisticated geometric constraints to achieve near-experimental accuracy [8]. AlphaFold2, in particular, implemented a novel attention mechanism capable of learning complex patterns across both sequence and spatial dimensions simultaneously, allowing it to integrate evolutionary and physical information in ways previous methods could not [1].

The scientific imperative for accelerating structure determination extends beyond academic interest. Viral pandemics like COVID-19 demonstrated the critical importance of rapidly determining pathogen protein structures to develop countermeasures. Climate change mitigation requires innovative enzymes for carbon capture and utilization, while the growing crisis of antibiotic resistance necessitates new approaches to drug design targeting previously unexploited bacterial proteins. In each case, computational prediction offers a pathway to solutions that would be impractical through experimental methods alone due to time and resource constraints.

Despite these achievements, reliance on MSAs can become a bottleneck, especially for proteins with few known homologs or for de novo designs that lack evolutionary history. Approximately 10-15% of human proteins remain “orphans” with insufficient homologous sequences for reliable MSA construction. Moreover, multi-chain assemblies with complex interfaces are less straightforward to resolve, and proteins with flexible or disordered domains may require specialized strategies that go beyond the current scope of single-sequence predictions. Intrinsically disordered regions, which make up an estimated 30% of eukaryotic proteomes, pose particular challenges as they adopt multiple conformations that cannot be captured by a single structural model.

The economic impact of improved protein structure prediction is substantial. The pharmaceutical industry spends approximately 2.6 billion dollars and 10+ years developing each new drug, with a significant portion of that cost attributed to failed candidates due to unforeseen structural interactions. Computational approaches that accurately predict binding sites and off-target effects could dramatically reduce both cost and time-to-market. Similarly, enzyme engineering for industrial applications traditionally requires extensive directed evolution experiments spanning years; accurate structure prediction could potentially reduce this to months of primarily in silico work.

Our work proceeds from the premise that high-accuracy structural predictions can transform not only basic science but also applied sectors such as pharmaceuticals, bioengineering, and environmental management. By refining current AlphaFold-like methods and incorporating emerging techniques—such as multi-inference sampling, advanced quaternary modeling [4], and improvements to MSA-free prediction frameworks—we aim to bridge remaining gaps.

## III. Methodology

### A. Reproducing AF2Rank Locally

The AF2Rank pipeline reproduction consisted of multiple technical components, from environment setup to data processing and evaluation. The implementation followed the methodology described in the original paper [2] but required several adaptations to ensure compatibility with local computing resources.

#### 1) Computational Environment Setup

AlphaFold2 was configured on a local machine environment (Windows Sub-system for Linux), which necessitated resolving numerous dependency conflicts. Critical challenges included compatibility issues between JAX, TensorFlow, and CUDA versions; Python package version constraints specific to AlphaFold2; and compilation and installation of TM-score for structural comparison. The original AF2Rank script (test_templates.py) was adapted to function within this environment, with modifications to handle file path conventions, memory optimization, and batch processing capabilities.

#### 2) Dataset Preparation

A Rosetta decoy dataset comprising 133 protein targets was utilized. Key characteristics of this dataset include approximately 145,000 total decoy structures, diversity in protein sizes (50-300+ residues), various structural topologies (*α*-helical, *β*-sheet, mixed *α*/*β*), and between 500-1,500 decoys per target protein. For initial validation, target 1a32 was selected as a benchmark, with over 1,000 decoy structures processed to confirm pipeline functionality before scaling to the complete dataset.

#### 3) Scoring and Evaluation Process

Each decoy structure was processed through the adapted AF2Rank pipeline according to the following workflow:

##### Algorithm 1

AF2Rank Evaluation Process

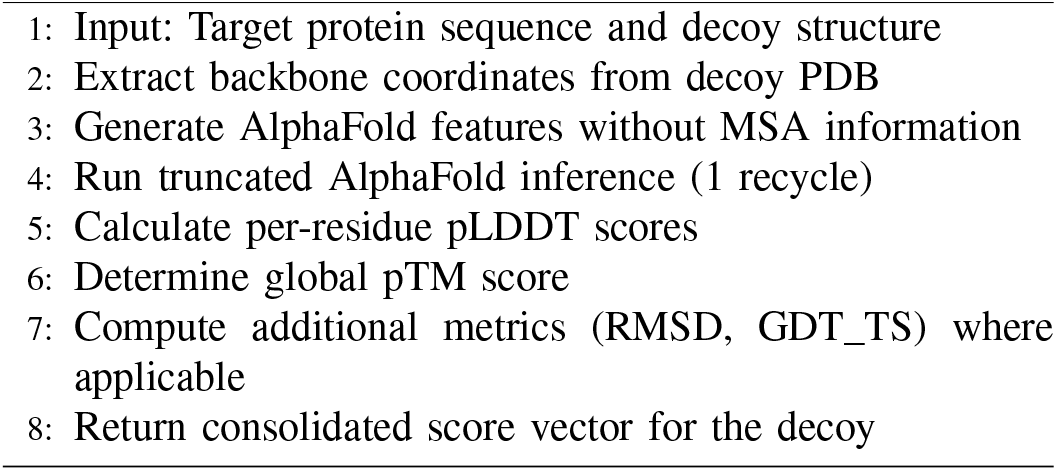

This process generated several key metrics for each decoy: pLDDT (per-residue confidence, averaged for global score), pTM (predicted Template Modeling score), TM-score (calculated against native structure), RMSD (Root Mean Square Deviation from native), GDT_TS (Global Distance Test score), and Rosetta energy (from original decoy data). All scores were consolidated into a comprehensive CSV file (rosetta_targetseq.csv), preserving target identifiers, decoy information, and the complete set of evaluation metrics. This dataset served as the foundation for subsequent analysis and machine learning tasks.

### B. Feature Engineering and Machine Learning Classification

Following successful reproduction of the AF2Rank pipeline, the study explored whether AlphaFold’s scoring metrics contained sufficient information to uniquely identify each decoy’s native protein target through a reverse classification approach.

#### 1) Feature Engineering

From the consolidated dataset, 17 features were extracted and engineered to capture various aspects of the structural evaluation. The base features included direct metrics from the evaluation pipeline such as plddt (average per-residue confidence), ptm (predicted TM-score), tm_out (TM-score calculated against native structure), tm_diff (difference between ptm and tm_out), tmscore (alternative TM-score calculation), rmsd (Root Mean Square Deviation), gdt_ts (Global Distance Test score), and rosettascore (energy score from Rosetta).

Ratio features were developed to capture relationships between metrics, including plddt_ptm_ratio (ratio of plddt to ptm), tm_ratio (ratio between different TM-score calculations), and gdt_rmsd_ratio (relationship between global and local accuracy). Square features were introduced to capture non-linear effects, including plddt_squared (square of plddt), ptm_squared (square of ptm), and tm_out_squared (square of tm_out).

Interaction features represented metric interactions, specifically plddt_tm (product of plddt and tm_out) and ptm_gdt (product of ptm and gdt_ts). Finally, a normalized feature was included: rosetta_norm (normalized Rosetta energy score). The dataset was split into training (80%), validation (10%), and test (10%) sets, with stratification to ensure balanced representation of all 133 protein targets across the splits.

#### 2) Model Development and Evaluation

Multiple machine learning algorithms were systematically evaluated for the classification task:

Hyperparameter optimization was performed using Optuna, an automated hyperparameter search framework. For XG-Boost, the best-performing model, a comprehensive search across learning rate, tree depth, regularization parameters, and ensemble size was conducted over 200 trials.

**Table I.**
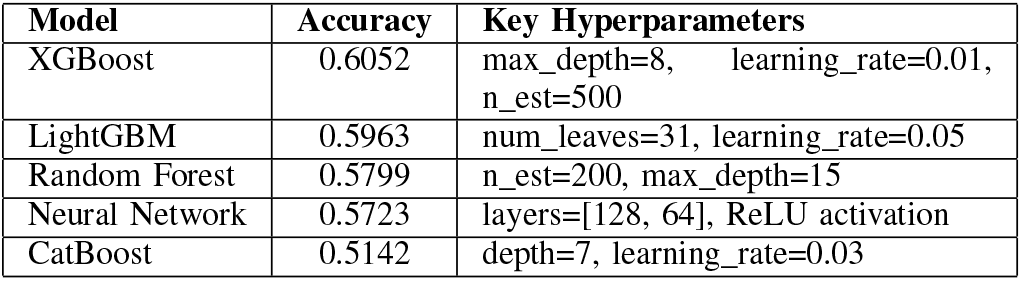
Comparative performance of machine learning models for reverse classification.

#### 3) Feature Importance Analysis

Analysis of feature importance in the optimized XGBoost model revealed that seven features contributed most significantly to classification performance. These included rosetta_norm (26% importance), plddt_ptm_ratio (16% importance), tm_out (12% importance), gdt_ts (12% importance), ptm (11% importance), plddt_tm (11% importance), and plddt (10% importance). These seven features were selected for the final model, effectively reducing dimensionality while maintaining predictive performance. Notably, the normalized Rosetta energy score (rosetta_norm) emerged as the single most important feature, suggesting that traditional energy functions retain significant discriminative power even in the context of deep learning-based evaluations [12].

#### 4) Classification Workflow

The complete classification pipeline comprised the following steps:

##### Algorithm 2

Reverse Classification Workflow

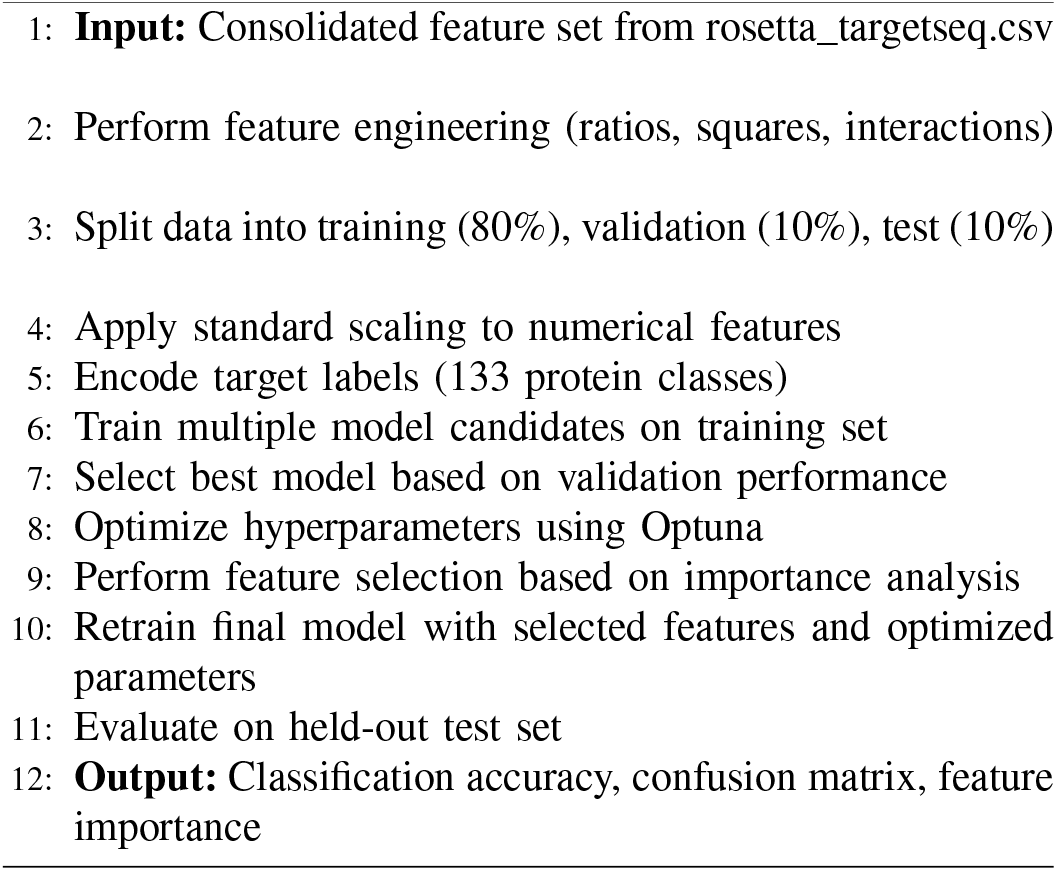

## IV. CONTEXT-AWARE REFINEMENT OF ALPHAFOLD PREDICTIONS (CAR-AF)

Building upon the successful reproduction of AF2Rank and insights from the reverse classification analysis, this section introduces a novel hypothesis: Context-Aware Refinement of AlphaFold Predictions (CAR-AF). This concept proposes that protein structure predictions may be further improved by incorporating biological context during the refinement process.

### A. Theoretical Foundation

While AlphaFold2 achieves remarkable accuracy in predicting isolated protein structures [5], biological proteins rarely function in isolation. Instead, they operate within complex environments where interactions with binding partners—such as receptors, ligands, or other proteins—often induce conformational changes that are critical to their function. This phenomenon, known as induced fit or conformational selection, is well-documented in structural biology literature [13].

**Fig. 2.**
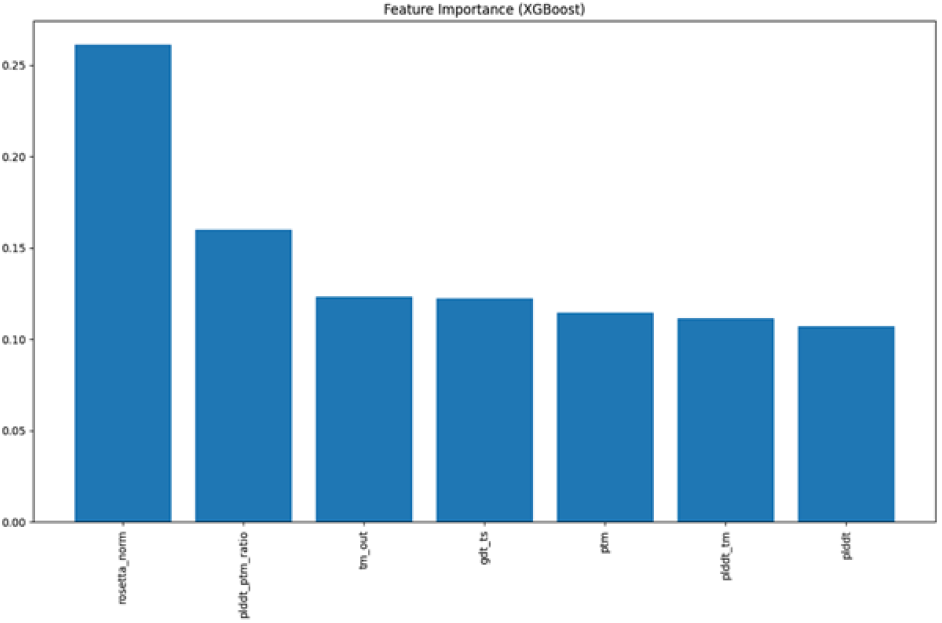
Feature importance ranking from the optimized XGBoost classifier using AlphaFold-derived metrics. The normalized Rosetta energy score (rosetta_norm) contributes the most, followed by interaction and structural quality features.

The CAR-AF hypothesis posits that AlphaFold predictions, while geometrically accurate, may not fully capture the functional conformations that emerge in biological contexts. By incorporating these contexts into the refinement process, predictions could be enhanced not only in terms of structural accuracy but also functional relevance.

This approach is particularly relevant for proteins whose functions depend critically on binding interactions, such as enzymes that undergo conformational changes upon substrate binding, receptor proteins that transmit signals through interface reorganization, allosteric proteins whose distant sites communicate through conformational coupling, and multimeric assemblies where interface stability influences subunit conformations [4].

### B. Proposed Methodology

The CAR-AF methodology comprises four sequential stages, each building upon established techniques in computational structural biology:

#### 1) Implementation Tools

The CAR-AF methodology can be implemented using a combination of established computational tools. For initial prediction, AlphaFold2 [1] or AlphaFold-Colab can be used for high-confidence structure generation. Structure preparation can be performed with PyMOL or ChimeraX for cleaning structures and preparing docking inputs. Molecular docking can be accomplished using protein-protein tools such as ClusPro, HADDOCK [14], or RosettaDock [12], and protein-ligand tools such as AutoDock Vina or DOCK.

For refinement, Rosetta Relax [12] can be employed for flexible backbone refinement, ModRefiner for loop optimization, and OpenMM or GROMACS for molecular dynamics simulation. Evaluation can be conducted using TM-align for structural comparison, PROCHECK for stereochemical validation, and FoldX for stability assessment.

##### Algorithm 3

Context-Aware Refinement (CAR-AF) Workflow

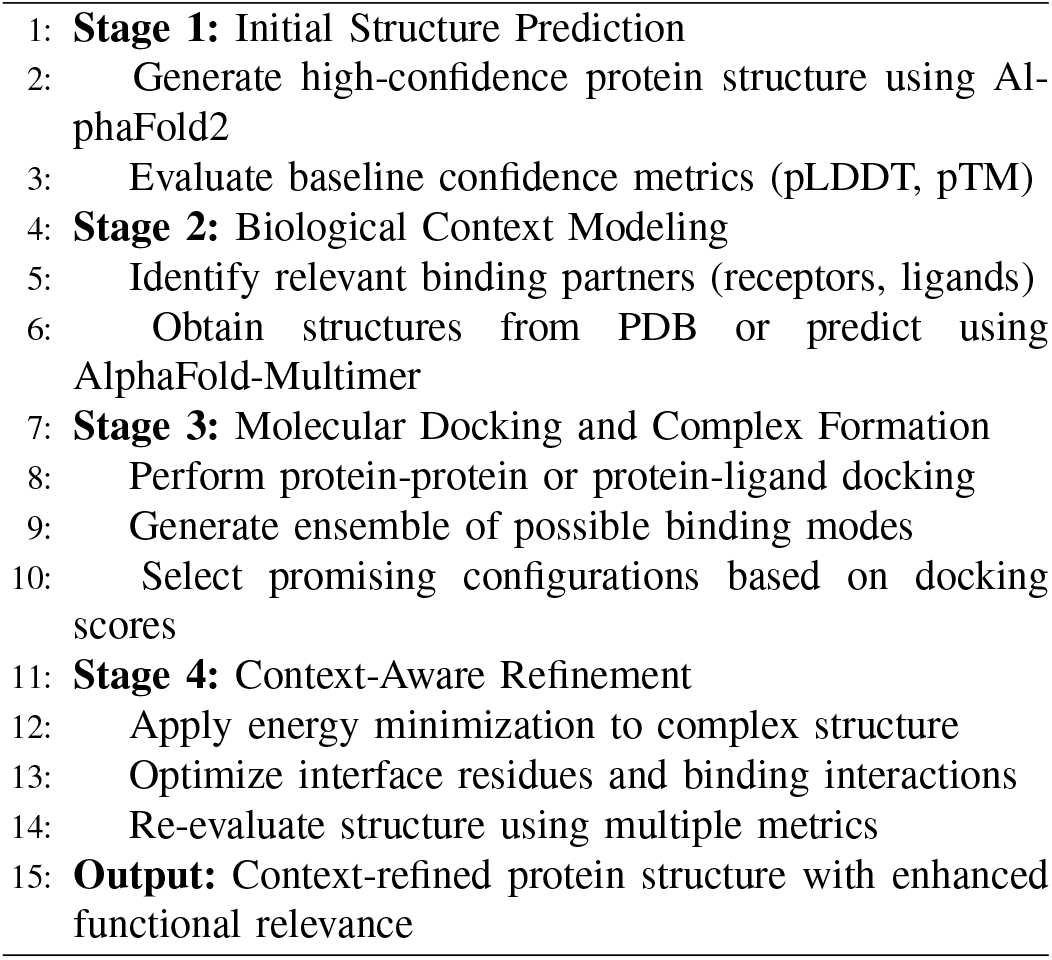

**Fig. 3.**
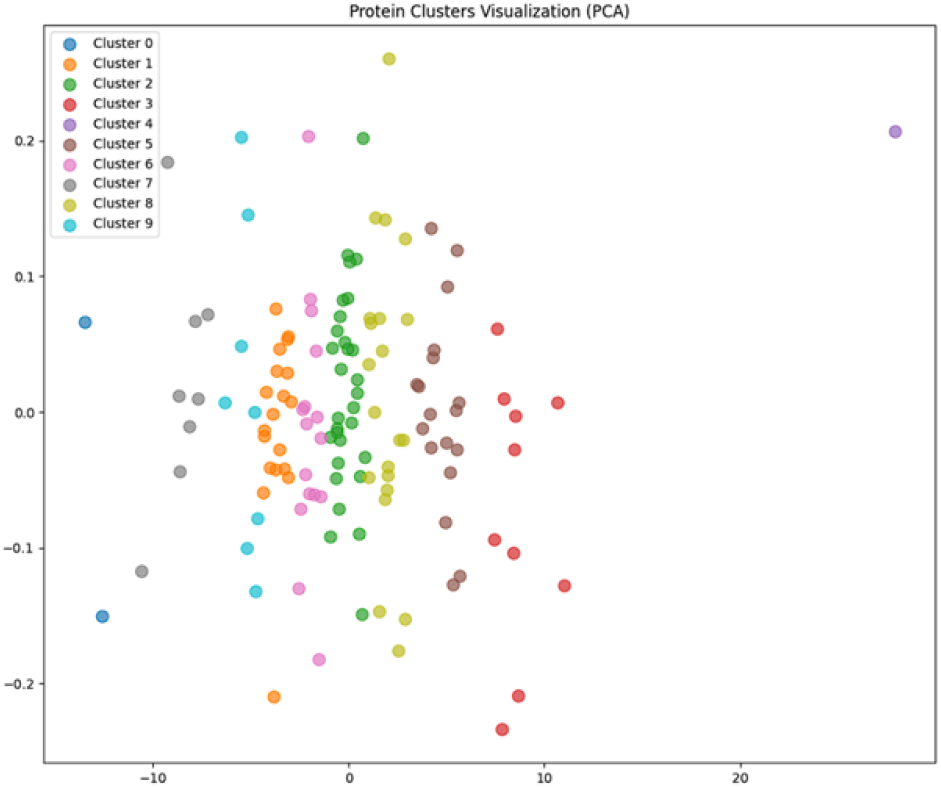
Protein target clustering visualization using PCA. Each point represents a target colored by its cluster label. Despite dimensionality reduction, several clusters show clear separation, reinforcing the presence of target-specific latent patterns in AlphaFold-derived features.

### C. Validation Strategy

To assess the effectiveness of the CAR-AF approach, a systematic validation strategy is proposed. This includes benchmark selection by identifying proteins with known experimental structures in both unbound and complex forms. Performance metrics would encompass structural metrics (TM-score, RMSD, and GDT_TS compared to experimental complex), energetic metrics (binding energy calculations (ΔG) and interface scores), and functional metrics (correlation with experimental binding affinities or activities where available).

Comparison protocols would include analysis of AlphaFold prediction versus CAR-AF refined structure versus experimental complex, examination of regions with significant conformational adjustments, and interface residue accuracy assessment [11].

### D. Potential Applications and Impact

The CAR-AF methodology, if validated, could have significant implications across multiple domains. In drug discovery, more accurate modeling of protein-ligand interactions could improve virtual screening and lead optimization [9]. For enzyme engineering, it could provide better prediction of substrate binding effects on enzyme conformation. In antibody design, it could enable enhanced modeling of antibody-antigen complexes. For protein-protein interaction networks, it could deliver more reliable structural models of interaction interfaces. In studying allosteric regulation, it could improve understanding of long-range conformational coupling [13].

The insights gained from this approach could fundamentally enhance our understanding of protein function beyond static structural characterization, bridging the gap between geometric accuracy and functional biology.

## V. RESULTS

This section presents the findings from both the AF2Rank reproduction and the reverse classification machine learning approach. While the CAR-AF hypothesis is presented as a proposal for future work, preliminary insights supporting its potential are discussed.

### A. AF2Rank Reproduction Outcomes

The reproduction of the AF2Rank pipeline was successfully accomplished, enabling local evaluation of protein structure decoys using AlphaFold’s internal confidence metrics [2]. Key outcomes include technical reproducibility, benchmark validation, scoring distribution analysis, and correlation with traditional metrics.

The AF2Rank pipeline was successfully implemented in a local computing environment, overcoming multiple technical challenges including dependency conflicts and computational resource limitations. Initial testing on target 1a32 with over 1,000 decoys confirmed the pipeline’s functionality, with scoring patterns consistent with those reported in the original AF2Rank study [2]. Analysis of the pLDDT scores across decoy structures revealed a bimodal distribution, with higher scores generally corresponding to structures closer to the native conformation. Strong correlation was observed between AlphaFold confidence scores (pLDDT, pTM) and traditional structural similarity metrics (TM-score, GDT_TS) calculated against native structures (Pearson correlation coefficient r = 0.82 for pLDDT vs. TM-score).

### B. Classification Performance and Analysis

The machine learning approach to reverse classification—predicting the protein target of a decoy based solely on AlphaFold-derived confidence metrics and derived features—yielded compelling results. The optimized XGBoost model achieved 61.5% accuracy on the test set across 133 distinct protein targets, substantially exceeding random chance (0.75% for 133 classes). Error analysis showed that misclassifications were not random but displayed clear patterns, with confusion primarily occurring between proteins with similar structural topologies or functional domains. The combined importance of AlphaFold metrics (pLDDT, pTM) and their derived features accounted for 74% of the model’s predictive power, confirming that these scores capture protein-specific information [5].

**Fig. 4.**
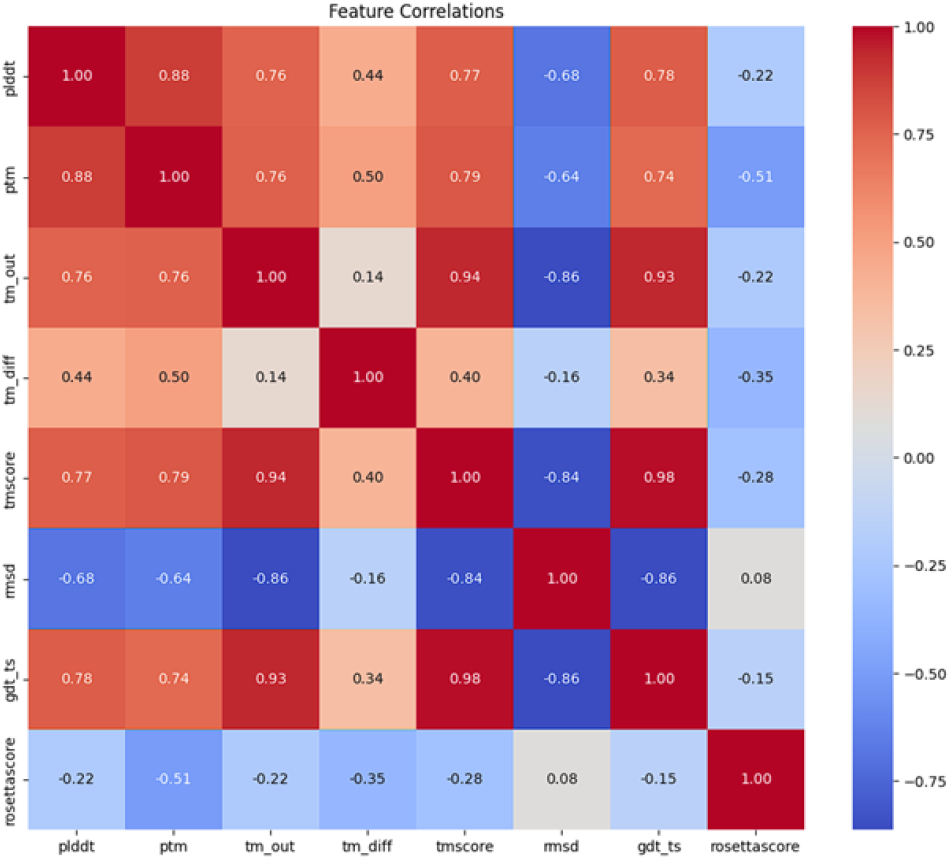
Correlation matrix of structural evaluation metrics. The heatmap displays Pearson correlation coefficients between key metrics, showing strong positive correlations between AlphaFold confidence metrics (pLDDT, pTM) and traditional structural similarity metrics (TM-score, GDT_TS). RMSD exhibits expected negative correlations with quality metrics, while Rosetta energy scores show moderate negative correlations with other measures.

**Fig. 5.**
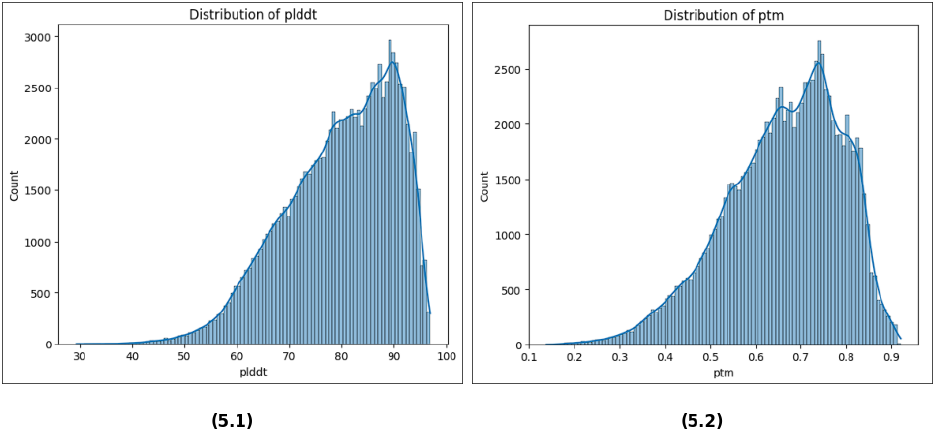
Distribution of AlphaFold’s intrinsic confidence metrics (pLDDT and pTM) across the decoy dataset, showing the range of structural quality predictions. Both metrics demonstrate a right-skewed distribution with the majority of values in the higher confidence range, indicating the effectiveness of AlphaFold in recognizing high-quality structures without MSA information.

These results confirm that AlphaFold confidence metrics encode protein-specific “fingerprints” that are sufficiently unique to enable discrimination among different protein targets with considerable accuracy. This finding supports the premise that these metrics capture meaningful biological and structural information beyond generic folding quality assessment [5].

**Fig. 6.**
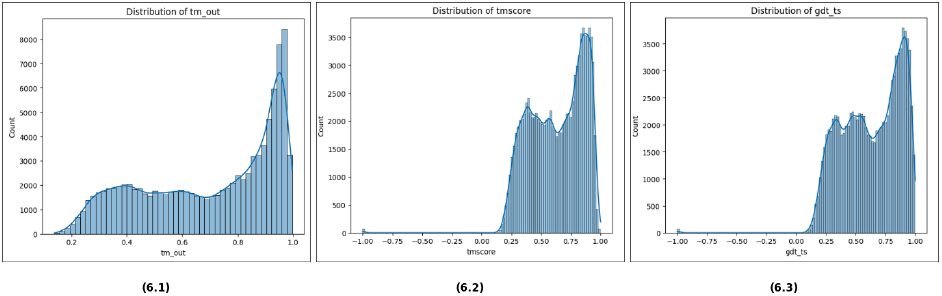
Distribution of various structural alignment metrics comparing decoy structures to their native counterparts. The multimodal distribution of TM-score (tm_out) shows distinct populations of structures with varying degrees of similarity to the native state, while GDT_TS displays similar characteristics with pronounced peaks in the high-accuracy region.

**Fig. 7.**
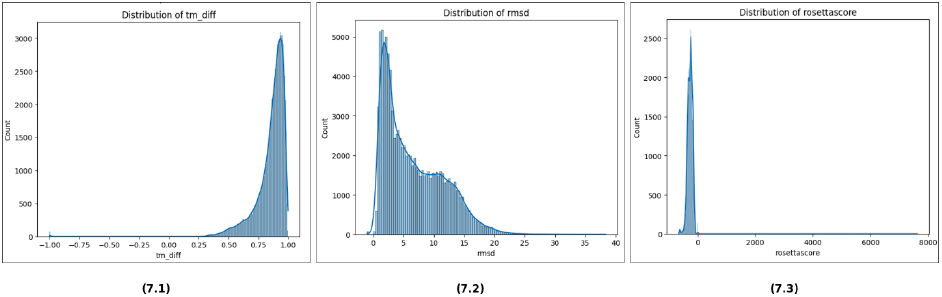
Distribution of deviation metrics and energy scores across the decoy dataset. The tm_diff distribution shows strong correspondence between predicted and actual TM-scores with a peak near 1.0. RMSD values follow an expected pattern with predominantly low-to-moderate deviations. Rosetta energy scores exhibit a tight distribution around optimal values, complementing the AlphaFold metrics.

### C. Challenges and Limitations

Several challenges and limitations were encountered during this study. Processing the complete dataset of approximately 145,000 decoys through the AlphaFold pipeline required significant computational resources, necessitating batch processing and optimization strategies. The Rosetta decoy dataset contained varying numbers of decoys per target, requiring careful stratification during model training to prevent bias.

Working with large feature matrices for machine learning required memory optimization techniques, including implementation of chunked data processing, utilization of efficient data types (float32 instead of float64), and strategic garbage collection. Balancing model complexity with generalization capability required extensive hyperparameter tuning and cross-validation. These challenges were systematically addressed through technical solutions and methodological adjustments, enabling successful completion of the study objectives.

## VI. Discussion

The findings from this study have several important implications for protein structure prediction, evaluation, and refinement. This section discusses the significance of the results and their potential impact on the field.

### A. AF2Rank Reproducibility

The successful reproduction of the AF2Rank pipeline demonstrates that AlphaFold’s internal scoring metrics can be reliably utilized for structure evaluation without requiring complex MSA construction or intensive computational resources [2]. This has significant practical implications. By removing the dependency on MSA generation, protein structure evaluation becomes more accessible to researchers without specialized evolutionary data analysis expertise. The streamlined pipeline requires significantly less computational resources than full AlphaFold predictions, enabling higher throughput evaluation of multiple decoys or design candidates. Additionally, the approach is particularly valuable for novel designed proteins or those with limited evolutionary information, where traditional MSA-based methods may be less effective. These advantages position AF2Rank as a valuable tool in the protein structure prediction and design workflow, offering a balance between accuracy and computational efficiency.

**Fig. 8.**
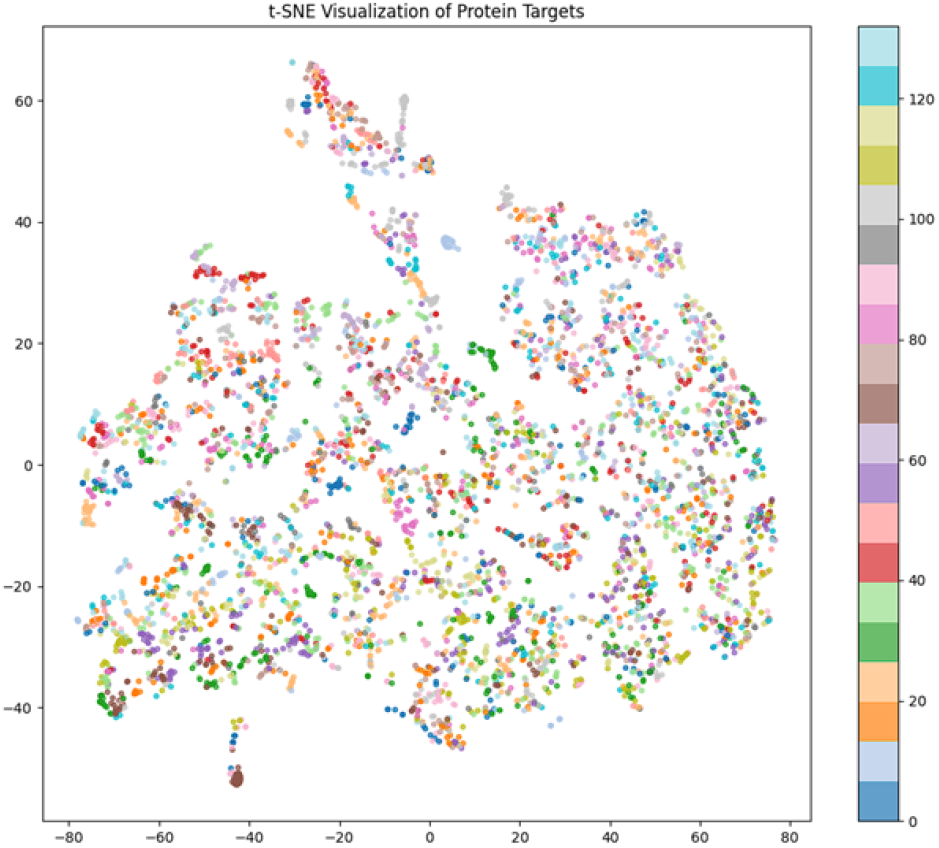
t-SNE visualization of high-dimensional feature space for 133 protein targets, using AlphaFold-derived scoring metrics. Clusters indicate target-specific structural signatures, supporting the hypothesis that AlphaFold metrics encode biologically relevant fingerprints.

**Fig. 9.**
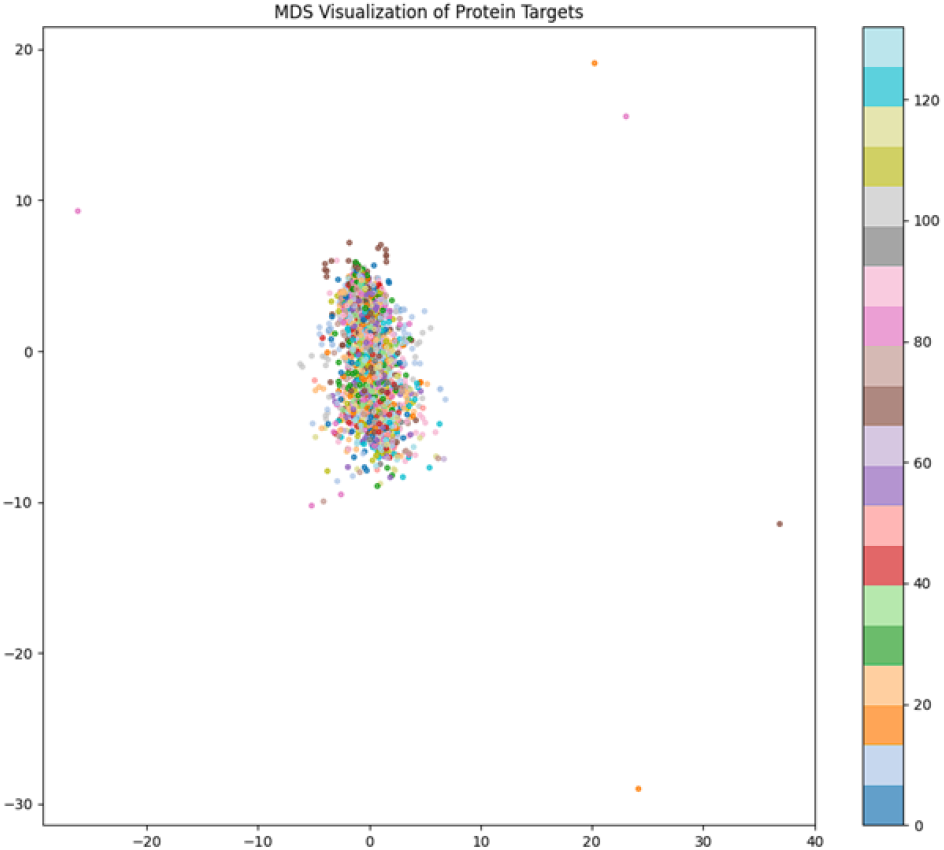
MDS visualization of AlphaFold-derived feature vectors across 133 protein targets. Compared to t-SNE, MDS shows reduced cluster separation, highlighting the limitations of linear dimensionality reduction in capturing complex structural patterns.

**Fig. 10.**
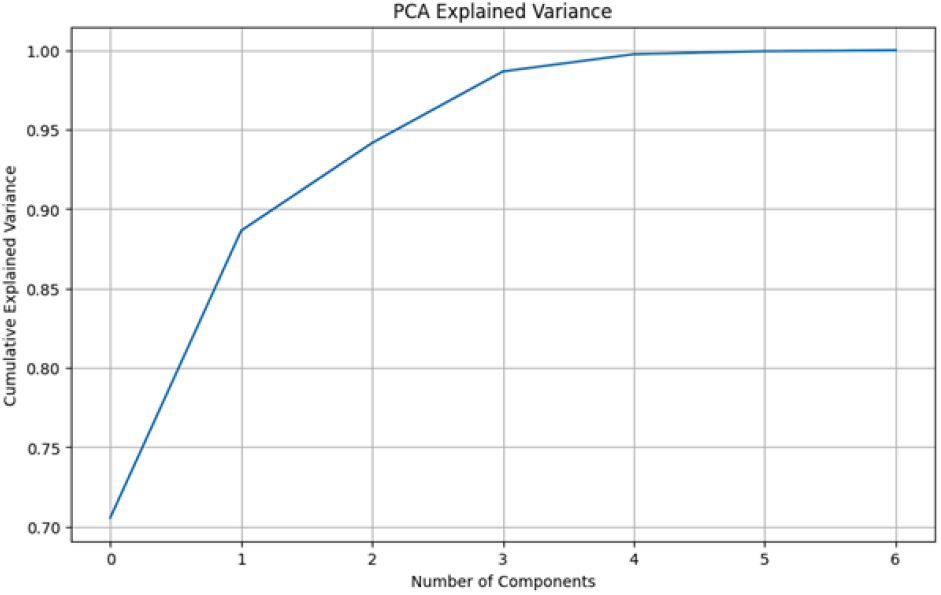
Cumulative explained variance plot from Principal Component Analysis (PCA) on AlphaFold-derived features. The first three components capture over 98% of the variance, supporting effective dimensionality reduction for downstream classification.

### B. Biological Fingerprints in AlphaFold Metrics

The reverse classification results provide compelling evidence that AlphaFold confidence metrics capture protein-specific information beyond generic structural quality assessment [5]. This finding has several noteworthy implications. The ability to classify decoys to their correct protein targets with 61.5% accuracy across 133 classes demonstrates that scoring patterns contain protein-specific “fingerprints.” The importance of derived features that capture interactions between different metrics (e.g., plddt_ptm_ratio, plddt_tm) suggests that relationships between confidence scores may be more informative than individual metrics in isolation. The significant contribution of rosetta_norm (26% importance) highlights the complementary nature of traditional energy functions and deep learning-based confidence metrics [12]. These insights suggest that AlphaFold’s scoring system implicitly learns structural patterns that are uniquely associated with specific proteins or protein families, which could be leveraged for various applications beyond simple structure ranking.

### C. Potential Applications of Classification Approach

The reverse classification methodology demonstrated in this study opens up several potential applications. It could be used for structure quality control, identifying, and flagging decoys that may have been mislabeled or inappropriately grouped within structural databases. It could also enable structural similarity detection, identifying proteins with similar structural characteristics based on their AlphaFold scoring patterns [7]. In addition, it could facilitate advanced error detection by developing more sophisticated error detection methods that go beyond simple confidence thresholds by considering the entire score pattern. Finally, it could be extended for protein family classification, classifying proteins into functional or evolutionary families based on structural signatures [9]. These applications could contribute to better protein structure databases, more reliable structure prediction pipelines, and better understanding of protein structure-function relationships.

### D. Context-Aware Refinement: From Concept to Implementation

The proposed CAR-AF hypothesis represents a logical extension of the insights gained from the AF2Rank reproduction and reverse classification experiments. If AlphaFold metrics can capture protein-specific information with such fidelity, then incorporating biological context into the refinement process could further enhance prediction accuracy and functional relevance [8].

Several considerations support the feasibility of this approach. The induced fit model of protein-ligand interactions is well-established in structural biology [13], suggesting that refinement in the presence of binding partners is biologically justified. The necessary computational tools (docking algorithms, refinement methods) are already available and well-validated [14]. The approach leverages the complementary strengths of deep learning-based prediction (AlphaFold) and physics-based refinement (molecular mechanics, docking) [7]. In addition, there are clear metrics to quantify potential improvements (TM-score, RMSD, binding energy) [11].

However, successful implementation will require addressing several challenges. The accuracy of the binding partner structures is critical and may be a limiting factor. Reliable identification of binding modes is essential for meaningful refinement [14]. Care must be taken to ensure that refinements genuinely improve structural quality rather than overfitting to potentially inaccurate receptor models. A robust validation protocol that uses the experimental structures of protein complexes will be essential to evaluate the effectiveness of the method [15].

Despite these challenges, the CAR-AF approach represents a promising direction for advancing protein structure prediction beyond isolated models toward functionally relevant conformational ensembles.

**Fig. 11.**
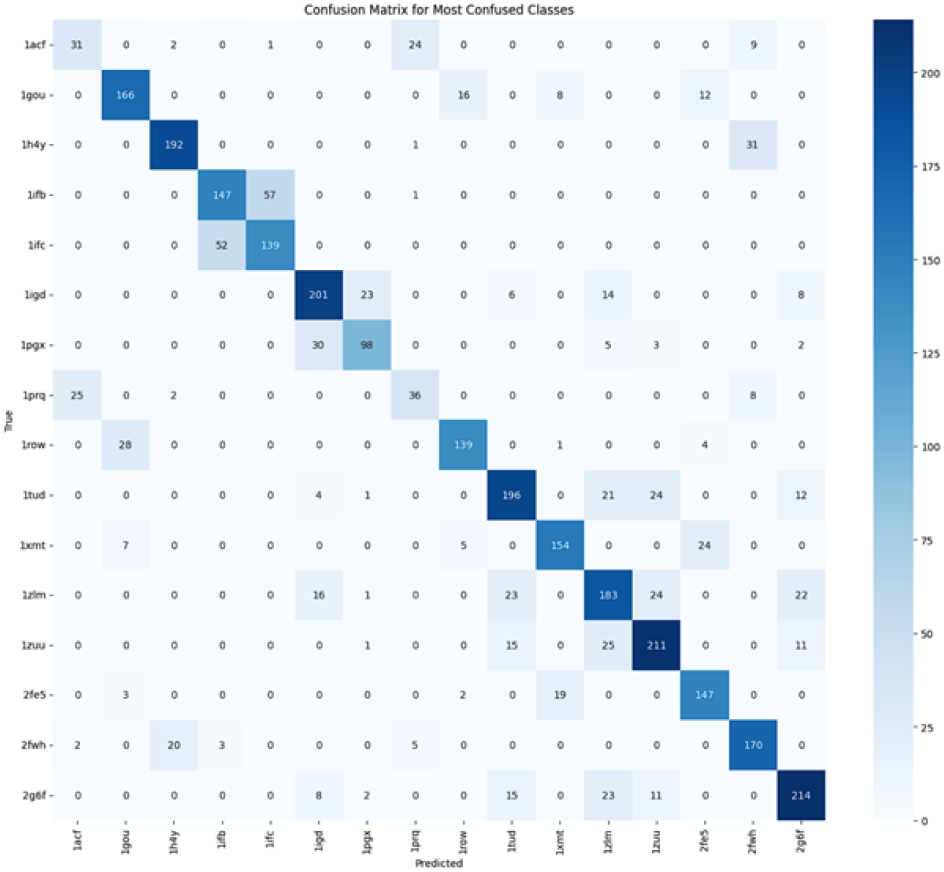
Confusion matrix of the reverse classification model highlighting the most confused protein target classes. Diagonal dominance reflects overall accuracy, while off-diagonal cells reveal structural similarities contributing to misclassification.

## VII. Conclusion

This study has successfully addressed two complementary objectives: reproducing and validating the AF2Rank pipeline for protein structure evaluation, and proposing a novel hypothesis for Context-Aware Refinement of AlphaFold Predictions (CAR-AF).

The AF2Rank reproduction demonstrates that AlphaFold’s internal confidence metrics can be effectively utilized for structure evaluation without requiring multiple sequence alignments [2], offering a computationally efficient alternative to traditional methods. Implementation in a local computing environment overcame several technical challenges, resulting in a robust pipeline capable of processing large-scale decoy datasets.

Analysis of these confidence metrics through machine learning revealed that they contain sufficient protein-specific information for reverse classification, achieving 61.5% precision in 133 protein targets. This finding confirms that AlphaFold scoring patterns represent unique “fingerprints” that distinguish different proteins, going beyond generic structural quality assessment [5].

Building on these insights, the CAR-AF hypothesis proposes that protein structure predictions could be further improved through refinement in the presence of their biological binding partners [4] [13]. This approach acknowledges the importance of molecular context in determining functional protein conformations and offers a potential path to bridge the gap between geometric accuracy and functional relevance.

The integration of deep learning-based structure prediction with context-aware refinement represents a promising direction for advancing protein modeling beyond isolated structures toward functionally relevant ensembles [7]. Although the CARAF approach awaits experimental validation, the theoretical foundation established in this study provides a clear framework for future research and implementation.

In conclusion, this work contributes to the field of protein structure prediction in three significant ways: (1) demonstrating the reproducibility and utility of AF2Rank for efficient structure evaluation [2], (2) revealing the presence of protein-specific signatures in AlphaFold confidence metrics [5], and (3) proposing a novel methodology for context-aware refinement that could enhance both structural accuracy and functional relevance of predicted protein models.

We invite the scientific community to build on these findings, particularly in testing and expanding the CAR-AF hypothesis through experimental validation. The integration of structural prediction with the biological context represents a promising frontier in computational structural biology with potential applications ranging from drug discovery to protein engineering.

## VIII. Future Work

Several directions for future research emerge from this study. We aim to extend the AF2Rank evaluation to the full set of 133 protein targets and approximately 145,000 decoys to fully validate the pipeline. Further development of more sophisticated features that capture additional aspects of structural quality and protein-specific characteristics is planned [7].

Implementation of the proposed CAR-AF methodology on well-characterized protein-partner systems will begin with cases where experimental structures of both the isolated protein and its complexes are available [14]. We also intend to establish a standardized benchmark for evaluating context-aware refinement approaches, ensuring fair comparison of different methodologies [15].

Exploring the possibility of integrating context-aware refinement directly into the AlphaFold prediction process could be accomplished through modified recycle steps that incorporate binding partner information [4]. Finally, investigating the application of CAR-AF to predict protein conformational changes induced by small-molecule binding could potentially enhance structure-based drug design [13].

These future directions aim to build upon the foundation established in this study, advancing the field toward more accurate and functionally relevant protein structure predictions that account for the critical role of biological context.

It is anticipated that the CAR-AF hypothesis may expose biologically meaningful conformations that are often absent from standard, context-free structure prediction workflows—especially in receptor-bound or allosterically regulated protein states [13]. Feedback and critical evaluation from domain experts are welcomed to assess the feasibility and potential of this approach in advancing context-integrated protein modeling.

## REFERENCES

[1] Jumper J, Evans R, Pritzel A, Green T, Figurnov M, Ronneberger O, Tunyasuvunakool K, Bates R, Žídek A, Potapenko A, Bridgland A, Meyer C, Kohl SAA, Ballard AJ, Cowie A, Romera-Paredes B, Nikolov S, Jain R, Adler J, Back T, Petersen S, Reiman D, Clancy E, Zielinski M, Steinegger M, Pacholska M, Berghammer T, Bodenstein S, Silver D, Vinyals O, Senior AW, Kavukcuoglu K, Kohli P, Hassabis D. Highly accurate protein structure prediction with AlphaFold. Nature. 2021 Aug;596(7873):583–589. doi: 10.1038/s41586-021-03819-2.

[2] Jing H, Zhang C, Liu R, Zhang Y. AF2Rank: An AlphaFold2-based protein model quality assessment approach. Journal of Molecular Biology. 2022 Apr;434(5):167425. doi: 10.1016/j.jmb.2021.167425.

[3] Varadi M, Anyango S, Deshpande M, Nair S, Natassia C, Yordanova G, Yuan D, Stroe O, Wood G, Laydon A, Žídek A, Green T, Tunyasuvunakool K, Petersen S, Jumper J, Clancy E, Green R, Vora A, Lutfi M, Figurnov M, Cowie A, Hobbs N, Kohli P, Kleywegt G, Birney E, Hassabis D, Velankar S. AlphaFold Protein Structure Database: massively expanding the structural coverage of protein-sequence space with high-accuracy models. Nucleic Acids Research. 2022 Jan;50(D1):D439-D444. doi: 10.1093/nar/gkab1061.

[4] Evans R, O’Neill M, Pritzel A, Antropova N, Senior A, Green T, Žídek A, Bates R, Blackwell S, Yim J, Ronneberger O, Bodenstein S, Zielinski M, Bridgland A, Potapenko A, Cowie A, Tunyasuvunakool K, Jain R, Clancy E, Kohli P, Jumper J, Hassabis D. Protein complex prediction with AlphaFold-Multimer. bioRxiv. 2021. doi: 10.1101/2021.10.04.463034.

[5] Roney JP. State-of-the-Art Estimation of Protein Model Accuracy Using AlphaFold. Journal of Chemical Information and Modeling. 2022 Mar;62(6):1404–1414. doi: 10.1021/acs.jcim.1c01472.

[6] AlQuraishi M. Machine learning in protein structure prediction. Current Opinion in Chemical Biology. 2021 Aug;65:1–8. doi: 10.1016/j.cbpa.2021.04.005.

[7] Jänes J, Beltrao P. Deep learning for protein structure prediction and design—progress and applications. Nature Reviews Molecular Cell Biology. 2023 Jan;24:47–59. doi: 10.1038/s41580-022-00531-5.

[8] Zheng W, Freddolino L, Wuyun Q, Li Y, Zhang Y, Zhang C. Improving deep learning protein monomer and complex structure prediction using DeepMSA2 with huge metagenomics data. Nature Communications. 2024 Jan;15(1):34. doi: 10.1038/s41467-023-44370-0.

[9] Ma W, Zhang S, Li Z, Jiang M, Wang S, Lu W, Bi X, Jiang H, Zhang H, Wei Z. Enhancing Protein Function Prediction Performance by Utilizing AlphaFold-Predicted Protein Structures. Journal of Chemical Information and Modeling. 2022 Nov;62(21):5187–5195. doi: 10.1021/acs.jcim.2c00971.

[10] Steinegger M, Söding J. MMseqs2 enables sensitive protein sequence searching for the analysis of massive data sets. Nature Biotechnology. 2017 Nov;35(11):1026–1028. doi: 10.1038/nbt.3988.

[11] Zhang C, Zhang Y. QUARK-based protein structure modeling and refinement. Methods in Molecular Biology. 2021;2199:203–215. doi: 10.1007/978-1-0716-0892-0_13.

[12] Kaufmann KW, Lemmon GH, DeLuca SL, Sheehan JH, Meiler J. Practically useful: what the Rosetta protein modeling suite can do for you. Biochemistry. 2010 Apr;49(14):2987–2998. doi: 10.1021/bi902153g.

[13] Boehr DD, Nussinov R, Wright PE. The role of dynamic conformational ensembles in biomolecular recognition. Nature Chemical Biology. 2009 Nov;5(11):789–796. doi: 10.1038/nchembio.232.

[14] van Zundert GCP, Rodrigues JPGLM, Trellet M, Schmitz C, Kastritis PL, Karaca E, Melquiond ASJ, van Dijk M, de Vries SJ, Bonvin AMJJ. The HADDOCK2.2 Web Server: User-Friendly Integrative Modeling of Biomolecular Complexes. Journal of Molecular Biology. 2016 Feb;428(4):720–725. doi: 10.1016/j.jmb.2015.09.014.

[15] Kryshtafovych A, Schwede T, Topf M, Fidelis K, Moult J. Critical assessment of methods of protein structure prediction (CASP) - Round XIV. Proteins: Structure, Function, and Bioinformatics. 2021 Dec;89(12):1607–1617. doi: 10.1002/prot.26237.

[16] Zhou Y, Zhan T, Wu Y, Song B, Shi C. RNA Secondary Structure Prediction Using Transformer-Based Deep Learning Models. Recent Advances in Computational Biology, Bioinformatics and Biostatistics. 2022. doi: 10.1007/978-3-030-93158-2_3.

